# HIV-1 Rev interacts with HERV-K RcREs present in the human genome and promotes export of unspliced HERV-K proviral RNA

**DOI:** 10.1101/822619

**Authors:** Laurie R. Gray, Rachel E. Jackson, Patrick E. H. Jackson, Stefan Bekiranov, David Rekosh, Marie-Louise Hammarskjöld

**Author notes:** **Corresponding Author** Marie-Louise Hammarskjold.

## Abstract

**Background:** The HERV-K (HML-2) viruses are the youngest of the human endogenous retroviruses. They are present as several almost complete proviral copies and numerous fragments in the human genome. Many HERV-K proviruses express a regulatory protein Rec, which binds to an element present in HERV-K mRNAs called the RcRE. This interaction is necessary for the nucleo-cytoplasmic export and expression of HERV-K mRNAs that retain introns and plays a role analogous to that of Rev and the RRE in HIV replication. There are over 900 HERV-K RcREs distributed throughout the human genome. Thus, it was of interest to determine if Rev could functionally interact with selected RcRE elements that map either to HERV-K proviruses or human gene regions. This interaction would have the potential to alter the expression of both HERV-K mRNAs and cellular mRNAs during HIV-1 infection.

**Results:** In this study we employed a combination of RNAseq, bioinformatics and cell-based functional assays. Potential RcREs were identified through a number of bioinformatic approaches. They were then tested for their ability to promote export and translation of a reporter mRNA with a retained intron in conjunction with Rev or Rec. Some of the selected elements functioned well with either Rev, Rec or both, whereas some showed little or no function. Rev function on individual RcREs varied and was also dependent on the Rev sequence. We also performed RNAseq on total and cytoplasmic RNA isolated from SupT1 cells expressing HIV Rev, with or without Tat, or HERV-K Rec. Proviral mRNA from three HERV-K loci (4p16.1b, 22q11.23 and most significantly 3q12.3) accumulated in the cytoplasm in the presence of Rev or Tat and Rev, but not Rec. Consistent with this, the 3’ RcRE from 3q12.3 functioned well with HIV-Rev in our reporter assay. In contrast, this RcRE showed little or no function with Rec.

**Conclusions:** The HIV Rev protein can functionally interact with many RcREs present in the human genome, depending on the RcRE sequence, as well as the Rev sequence. This leads to export of some of the HERV-K proviral mRNAs and also has the potential to change the expression of non-viral genes.

## Background

With the recent advances in DNA sequencing, it is now realized that human endogenous retroviruses (HERVs) constitute about 8% of the human genome. This is a result of retroviral integration after infection of the germ line. More than 200,000 copies of HERVs have been identified, making up more than thirty distinct groups (1, 2). The majority of these proviruses are silenced through chromatin modifications and the accumulation of multiple point mutations, deletions, or insertions in their open reading frames (ORFs) or regulatory elements (3–5). Nevertheless, multiple endogenous retroviral copies are still almost complete and many fragmented HERV sequences provide functional *cis*-acting signals, such as promoters, enhancers, alternative splice sites, and poly-A signals that can alter human gene expression (6–8).

The youngest HERV family, HERV-K(HML-2), hereafter referred to as HERV-K, is thought to have integrated into the germ line between 200,000 and 5 million years ago (9, 10). HERV-K viruses are named for their utilization of a lysine tRNA to prime reverse transcription and the Human MMTV-Like (HML-2) designation refers to the fact that these viruses are similar to mouse mammary tumor virus (MMTV) (11). The copies of these viruses in the human genome have accumulated the fewest number of mutations among the HERVs and several have retained ORFs (8, 11). Two different types of HML-2 proviruses have been identified, based on their potential to produce different viral regulatory proteins. Type I HERV-K viruses encode a protein named Np9, while type II viruses encode the regulatory protein Rec. Type I viruses are the result of a 292bp deletion in the genome that shifts the *rec* ORF into another reading frame in a second coding exon, making the sequence of Np9 almost completely different from Rec. The function of Np9 remains unknown, although some studies suggest that it may have oncogenic potential (11–13). Rec functions to facilitate the export of incompletely spliced HERV-K mRNAs from the nucleus to the cytoplasm (see below).

To date, over 90 copies of intact or partially intact HERV-K proviruses have been identified. Although infectious HERV-K molecular clones have been created in the laboratory (14–16), none of the sequences found in the human genome appear to produce a replicating, full-length integration-competent virus, despite the fact that some have complete open reading frames and two Long Terminal Repeats (LTRs) (14). Furthermore, many of these viruses are polymorphic in the human population, and new copies are still being discovered, so it remains possible that some of them are still capable of active retro-transposition (17, 18). In addition to the proviral copies, more than 900 solo HERV-K LTRs, which have arisen from recombination events, are also present (14). Many of these solo LTRs are located within intergenic regions of the genome, but they are also found within protein coding human genes (19). Since these LTRs contain sequences that can function as regulatory elements of transcription, they have the potential to influence cellular gene expression within a gene or up to 200kb either upstream or downstream of a neighboring gene (20).

The LTR of the HERV-K provirus also contains a sequence element, called the RcRE, that is present near the 3’ end of all of the HERV-K mRNAs. The interaction of this element with the HERV-K Rec protein forms a complex, together with Crm-1 and Ran-GTP (21–23), that promotes nuclear export and expression of the viral mRNAs that retain introns (24). Complex exogenous retroviruses, such as HIV (25, 26), HTLV (27, 28), JSRV (29) and MMTV (30) also utilize similar mechanisms. For example, the HIV viral Rev protein acts in conjunction with the Rev Responsive Element (RRE) (25, 26) to perform this task. Early studies, reported about 15 years ago, examining putative functional RcREs from unknown locations in the genome, showed that the HIV Rev protein could promote export and expression of mRNA with retained introns from reporter constructs containing these RcREs. It was also reported that although Rev functioned to some extent with RcREs, the prototype Rec proteins failed to function in conjunction with the HIV-1 RRE (23, 31). As discussed above, since hundreds of HERV-K LTRs are widely distributed throughout the human genome, potentially functional RcREs are also present in many human RNA transcripts. This raises the distinct possibility that when Rev is expressed during HIV infection, mRNA isoforms with retained introns, which would normally be found only in the nucleus, could be transported to the cytoplasm to alter the transcriptome. They could potentially also be translated and alter the proteome.

Several recent studies have examined the effect of HIV infection on host cell gene expression. A recent RNAseq study reported that HIV infection in primary CD4 T-cells led to accumulation of RNAs with a significant amount of intron retention in ribosomal protein and other genes (32). The sequence analysis also detected induction of novel RNAs from retrotransposons and endogenous retroviruses. This study did not provide an explanation for the observed increase in these mRNA species, although an effect of expression of Rev was discussed as a potential explanation.

Several reports have also demonstrated expression of HERV-K mRNAs and proteins from proviruses in HIV-1 infected patients (33), as well as in cultured cells infected with HIV (34). HERV-K virus-like particles have also been detected in the plasma of HIV-1 infected individuals (35). However, in a recent report, increased HERV-K virion production in HIV-1 infected individuals was not detected, although there was increased transcription from some HERV-K LTRs (36). Other reports have suggested that HERV-K transcripts and proteins can be induced by the HIV-1 viral proteins Tat and Vif, as a result of activation of transcription from the HERV-K LTR (37, 38).

Based on these studies, we decided to investigate whether putative functional RcREs found within known human genes, as well as in a HERV-K provirus (HERV-K 3q12.3) that was shown to be active during HIV infection, could functionally respond to HIV Rev and HERV-K Rec proteins. The sequences were chosen based on their degree of homology to the prototypical RcREs that had been tested to date. Since our previous studies showed that Rev/RRE pairs from different HIV primary isolates displayed striking differences in functional activity (39), we also analyzed if different Rev proteins functioned differentially on different RcREs and how well the Revs worked in comparison to the prototypical Rec. Here we report that the selected HERV-K RcREs functioned with varying efficiencies in response to different HIV-1 Rev proteins and that some in fact functioned better with some of the Rev proteins than with Rec. We also show that the 3’LTR of HERV-K 3q12.3 functioned very well with Rev, while the 5’ LTR did not and that the prototypical Rec protein showed almost no function on either element. Furthermore, expression of NL4-3 Rev, but not Rec, in SupT1 cells resulted in a significant increase in cytoplasmic, unspliced mRNA from this HERV-K provirus, and to a lesser extent, in at least two other ones.

## Results

### Selection of genomic RcRE sequences for functional testing

We initially performed BLAST searches in human RNA databases using one of the previously analyzed RcREs (which was named prRcRE for prototype RcRE) (23). Surprisingly, no sequences were found that showed 100% identity with the prRcRE or in our subsequent searches of the human genome. Based on these searches, we selected four sequences that showed high sequence identity to the prRcRE, present in RNAs expressed from the following genes (percent identity with the prRcRE indicated in parenthesis): C11orf72 (98%), RP4linc (97%) cFLAR (95%) and U9linc (94%) (see Figure 1, top 5 lines). To determine if the selected RcREs could function with either Rev or Rec similarly to the prRcRE, the sequences were cloned into the pCMV-Gag-Pol reporter vector (40). This vector produces a HIV GagPol mRNA containing the coding region within an intron, similarly to the way it is arranged within the HIV provirus (40). Because of this, the mRNA cannot be exported from the nucleus to the cytoplasm and translated into protein without the insertion of a *cis*-acting element capable of binding to a *trans*-acting export factor. Gag and GagPol proteins are only produced in the presence of both a *cis*-acting element and a *trans*-acting export factor that can directly interact with it. Gag and GagPol proteins assemble into HIV particles and are released into the medium, where they can be easily measured by a p24 ELISA. This vector has been used by us in multiple previous publications to measure the activity of many *cis*-acting elements and *trans*-acting factors that mediate export of mRNA with retained introns (40–43).

**Figure 1.**
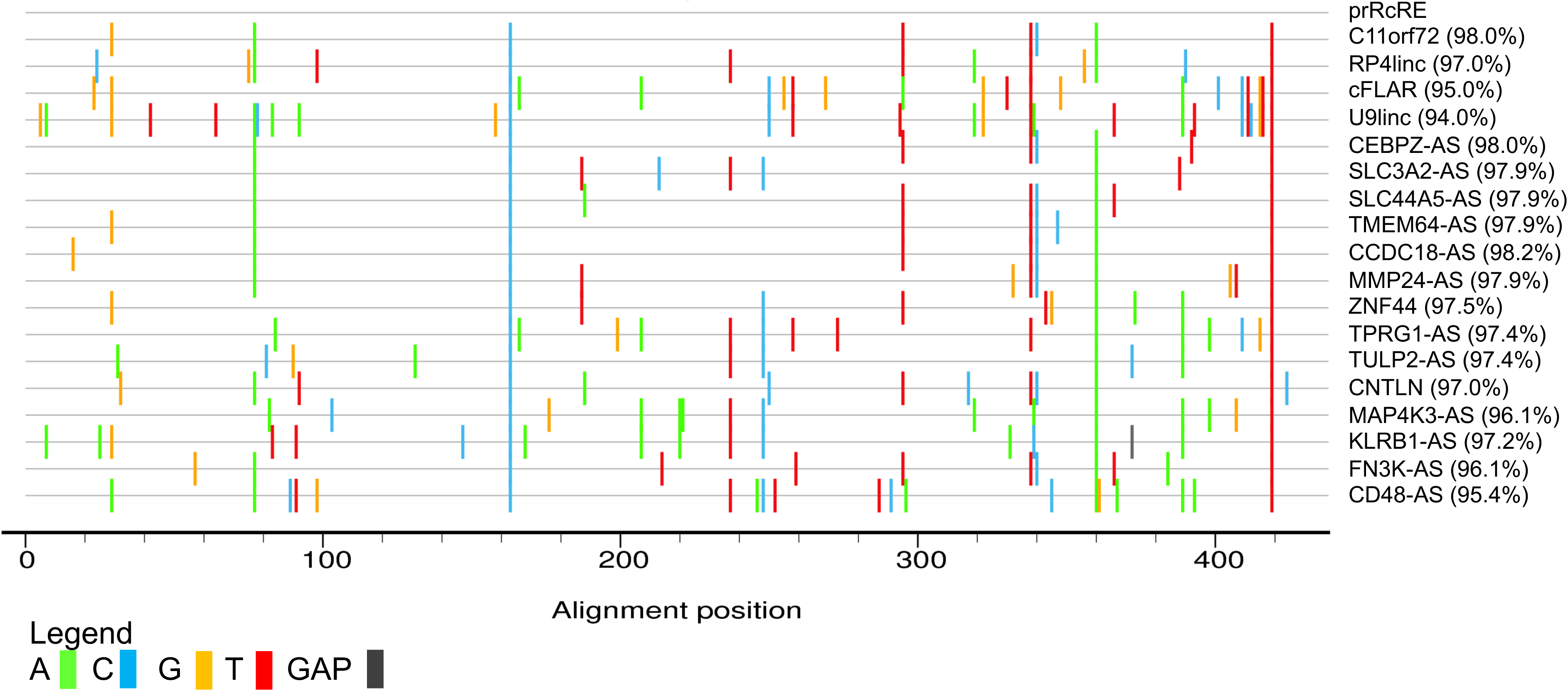
Highlighter plot of selected HERV-K RcREs. The sequences of the eighteen RcREs identified in this study that mapped to specific gene regions, were aligned to the prototypical RcRE. Mismatches are indicated by the colored lines. The sequences are listed according to the order in which they were tested in the results section. The results for the first four sequences are shown in Figure 2 and the results for the next 11 sequences are shown in Figure 4a and 4b. The remaining 3 sequences were not tested. Sequence identity of the selected RcREs to the prRcRE ranged from 94.0 to 98.2%.

After cloning of these elements into the vectors, they were transfected into 293T/17 cells together with plasmids expressing either HIV NL4-3 Rev or prRec (39, 43). The results of these experiments are shown in Figure 2. Without the co-expression of an export protein, most of the elements failed to elicit any p24 production, although low level activity was observed with RP4linc, cFLAR and U9linc. The element located within C11orf72 functioned significantly better than all of the others and both Rev and Rec functioned in conjunction with this element, although Rec showed higher activity than Rev (Figure 2A). The element located within the RP4linc gene also gave significant, albeit lower activity, with both Rev and Rec (Figure 2B), whereas the other two elements showed little change in functional activity in conjunction with either Rev or Rec. We also tested the GagPol vector containing the RRE, which confirmed previous results that indicated that Rec does not function in conjunction with the HIV-RRE (Figure 2C). In contrast, both NL4-3 Rev and Rec functioned on the prRcRE, consistent with previous data. These experiments indicate that some human mRNAs and lncRNAs contain elements that function as RcREs, capable of working with both Rec and Rev proteins. However, they also showed that even though many sequences show high sequence identity to functional RcREs, even a few nucleotide changes can cause significant activity differences.

**Figure 2.**
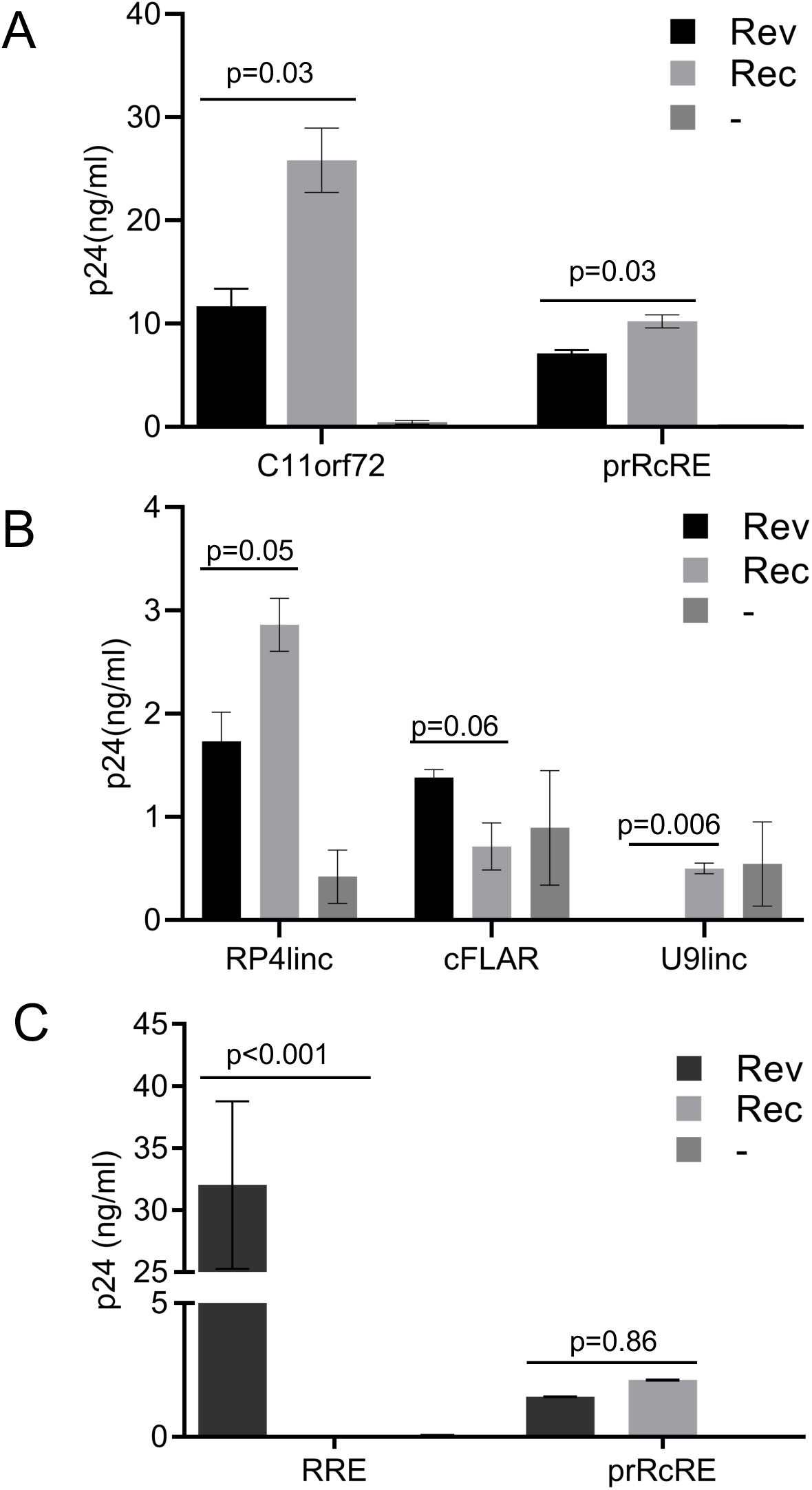
Functional activity of selected HERV-K RcREs in a transient transfection assay. The functional activities of the RcREs mapping to C11orf72, cFLAR, RP4linc and U9linc were measured by transfecting the reporter constructs containing each RcRE into 293T/17 cells together with either 50ng of prRev or prRec or no additional vector. As controls, reporter constructs containing the prRcRE and HIV-1 RRE were also tested. After 72 hours, cell supernatants were harvested and supernatant p24 was measured by ELISA. The error bars show the variation between two independent transfections performed at the same time. To determine p values, between Rev and Rec, a two-tailed student t-test was performed using GraphPad Prism 8. Note the different scales shown in A, B and C. The results shown in C are from an experiment performed on a different day than the results shown in A and B.

### Selection of RcRE sequences in the human genome mapping to annotated gene regions that are common to datasets from HIV-1 infected cells and cells expressing Rec proteins

Since our results demonstrated that primary sequence alone was not sufficient to identify whether a specific element was functional with Rev or Rec, we next employed a multi-step approach to increase the probability of finding functional elements. First, we performed a BLAST search with the prRcRE against the Ensembl DNA human genomic GRCh38.87 database (44). This resulted in a dataset of 198 sequences with greater than 94% sequence identity with the prRcRE. Within this dataset, 88 potential unique functional RcRE sequences were found to be present in 98 annotated gene regions, as some of these regions had duplicate or identical RcRE sequences. We then queried our dataset to find genes that had been previously shown to have altered expression after HIV-1 infection (32) and/or genes that were differentially expressed in human embryonic carcinoma cells (hECC) expressing exogenously added Rec (45) (Figure 3). Genes were considered differentially expressed in these datasets if the false discovery rate (FDR) adjusted p-values were <0.01. Three genes, (CEBPZ, SLC44A5 and SLC3A2) were common to all three datasets. A fourth annotated gene, TMEM64, was shared between our dataset and the Rec dataset, but was absent from the HIV dataset. The four potential RcREs were highly similar, showing 97-98% identity to the prRcRE (Figure 1). All of these elements are located within regions that have been designated as introns and are present in the anti-sense orientation relative to the transcription of the respective gene. The RcRE in the SLC44A5 gene region is located within the 3’ LTR of a partial HERV-K provirus, 1p31.1. The TMEM64 gene region contains two solo direct repeat LTRs (451 and 452) with identical RcREs.

**Figure 3.**
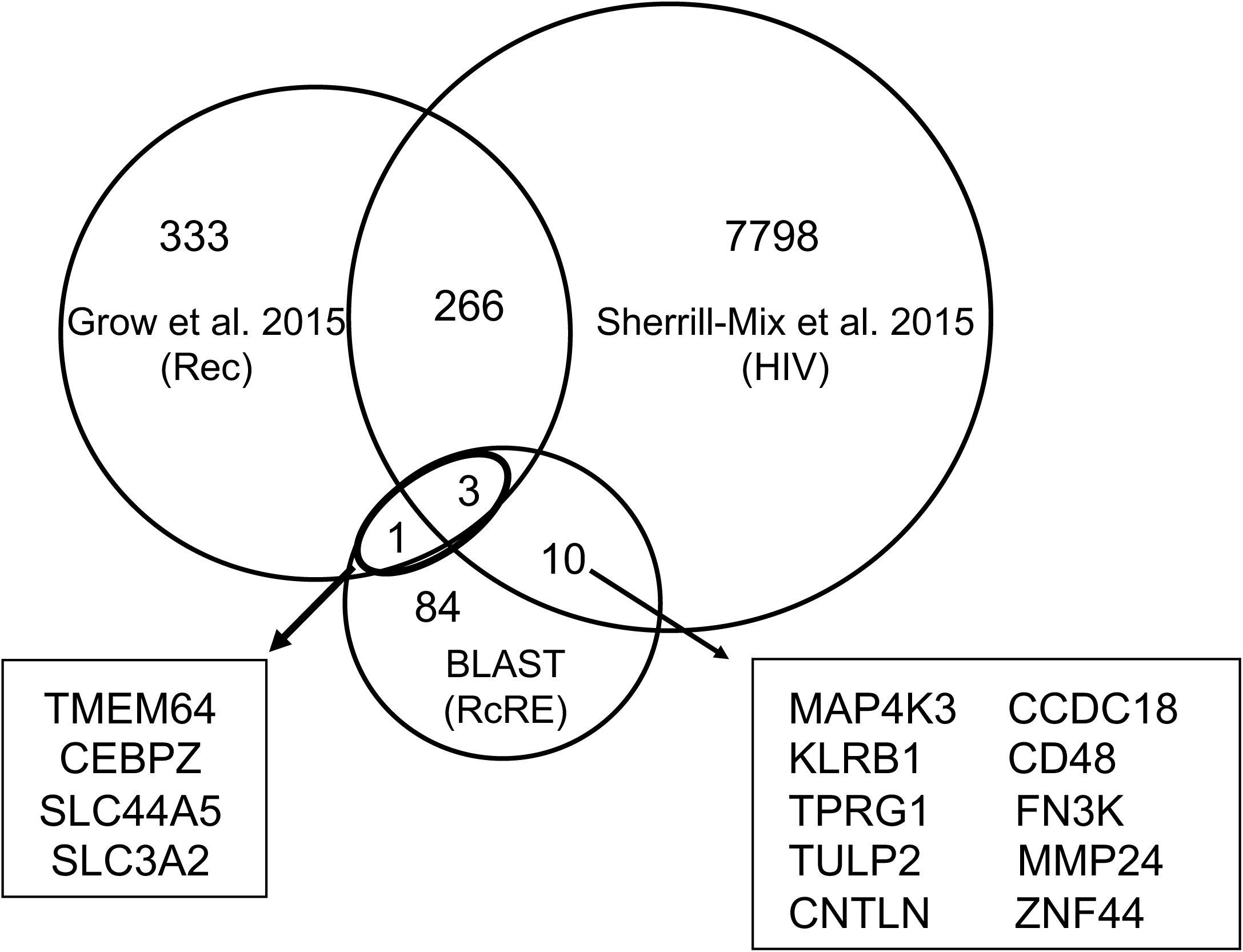
Venn diagram showing overlap between gene regions containing a RcRE and genes altered after HIV infection or differentially expressed in hECC in response to Rec. The dataset named RcRE contained sequences in gene regions that were highly similar to the HERV-K prRcRE. The dataset named HIV, contained a list of genes that were shown to be changed after HIV-1 infection (32). The dataset named Rec contained genes that were differentially expressed in human embryonic carcinoma cells (hECC) expressing HERV-K Rec (45). Three genes, CEBPZ, SLC44A5 and SLC3A2, were common to all three datasets. A fourth annotated gene, TMEM64, was shared between the Rec and RcRE datasets, but was absent from the HIV dataset. Ten additional annotated genes from the HIV dataset contained RcREs, however, these genes were absent from the Rec dataset. The RcREs located within CNTLN and TPRG1 are in the same transcriptional orientation as the gene, while the remaining RcREs are anti-sense to the transcriptional orientation of their associated gene.

We also identified genes that were common to only the RcRE similarity and HIV datasets. The ten identified genes, (MAP4K3, TPRG1, TULP2, CNTLN, CCDC18, MMP24, KLRB1, FN3K, ZNF44 and CD48) all contained elements located within introns that are similar (95.4-98.2%) to the prRcRE (Figure 1). Two of the genes, CNTLN and ZNF44, contained elements that are sense relative to the transcription of the gene. The remaining eight genes, (MAP4K3, TPRG1, TULP2, CCDC18, MMP24, KLRB1, FN3K and CD48) contained elements that are anti-sense to the transcription of the gene. The element in CD48 is located within the 3’ LTR of HERV-K provirus, 1q23.3. The locations of these elements are shown in Table 1 with the LTR annotation adopted from Subramanian et al (14).

**Table 1.** Summary of RcREs identified in this study. The RcRE sequences identified in this study are summarized by their Ensembl Gene ID, Hugo Gene Nomenclature Committee name (HGNC), chromosomal location in GRCh38.87/hg38, whether the sequence is sense or anti-sense to the gene transcription and HERV-K LTR5Hs annotation adopted from Subramanian et al. (14)

| Gene | HGNC | RcRE Location (sense*) | LTR # |
| --- | --- | --- | --- |
| ENSG00000272455 | RP4linc | chr1:1,410,022-1,410,454 | 1 |
| ENSG00000003402 | cFLAR | chr2:201,139,217-201,139,650 | 139 |
| 3q12.3 (K(II), ERVK-5) |  | chr3:101,700,268-101,700,695 | - |
| ENSG00000272462 | U9linc | chr6:25,999,448-25,999,878 | - |
| ENSG00000044459 | CNTLN | chr9:17,445,576-17,446,008 | 472 |
| ENSG00000184224 | C11orf72 | chr11:67,603,674-67,604,106 | 575 |
| ENSG00000197857 | ZNF44 | chr19:12,232,143-12,232,575 | 765 |
| Gene | HGNC | RcRE Location (anti-sense*) | LTR |
| ENSG00000137968 | SLC44A5 | chr1:75,377,301-75,377,733 | - |
| ENSG00000122483 | CCDC18 | chr1:93,277,102-93,277,534 | 40 |
| ENSG00000117091 | CD48 | chr1:160,691,001-160,691,433 | - |
| ENSG00000115816 | CEBPZ | chr2:37225,533-37,225,965 | 108 |
| ENSG00000011566 | MAP4K3 | chr2:39,321,376-39,321,808 | 111 |
| ENSG00000188001 | TPRG1 | chr3:189,251,690-189,252,123 | 208 |
| ENSG00000180694 | TMEM64 | chr8:90,684,247-90,684,679 | 451,452 |
| ENSG00000168003 | SLC3A2 | chr11:62,858,836-62,859,268 | 570 |
| ENSG00000111796 | KLRB1 | chr12:9,600,681-9,601,113 | 606 |
| ENSG00000167363 | FN3K | chr17:82,744,850-82,745,277 | 742 |
| ENSG00000104804 | TULP2 | chr19:48,889,851-48,890,283 | 809 |
| ENSG00000125966 | MMP24 | chr20:35,262,444-35,262,876 | 867 |

### Functional testing of potential RcRE elements in the identified gene regions

We decided to first test the four potential RcREs that were common to both the Rec dataset and our dataset. To do this, synthetic DNA fragments containing the RcRE sequences were cloned into the HIV-1-Gag-Pol reporter construct. The resulting plasmids were then transfected into 293T/17 cells together with either pCMV-Rev or pCMV-Rec. The results of these experiments demonstrated that all of these elements functioned well with both Rev and Rec (Figure 4A). Interestingly, the different RcREs responded differentially to Rev and Rec. In the case of the RcREs from CEBPZ, SLC44A5 and TMEM64, higher levels of p24 were obtained with Rec than with Rev. In contrast, the RcRE from the SLC3A2 gene region showed significantly higher levels of p24 expression with Rev compared to the levels obtained with Rec. SLC3A2 also showed differential expression after infection with HIV.

**Figure 4.**
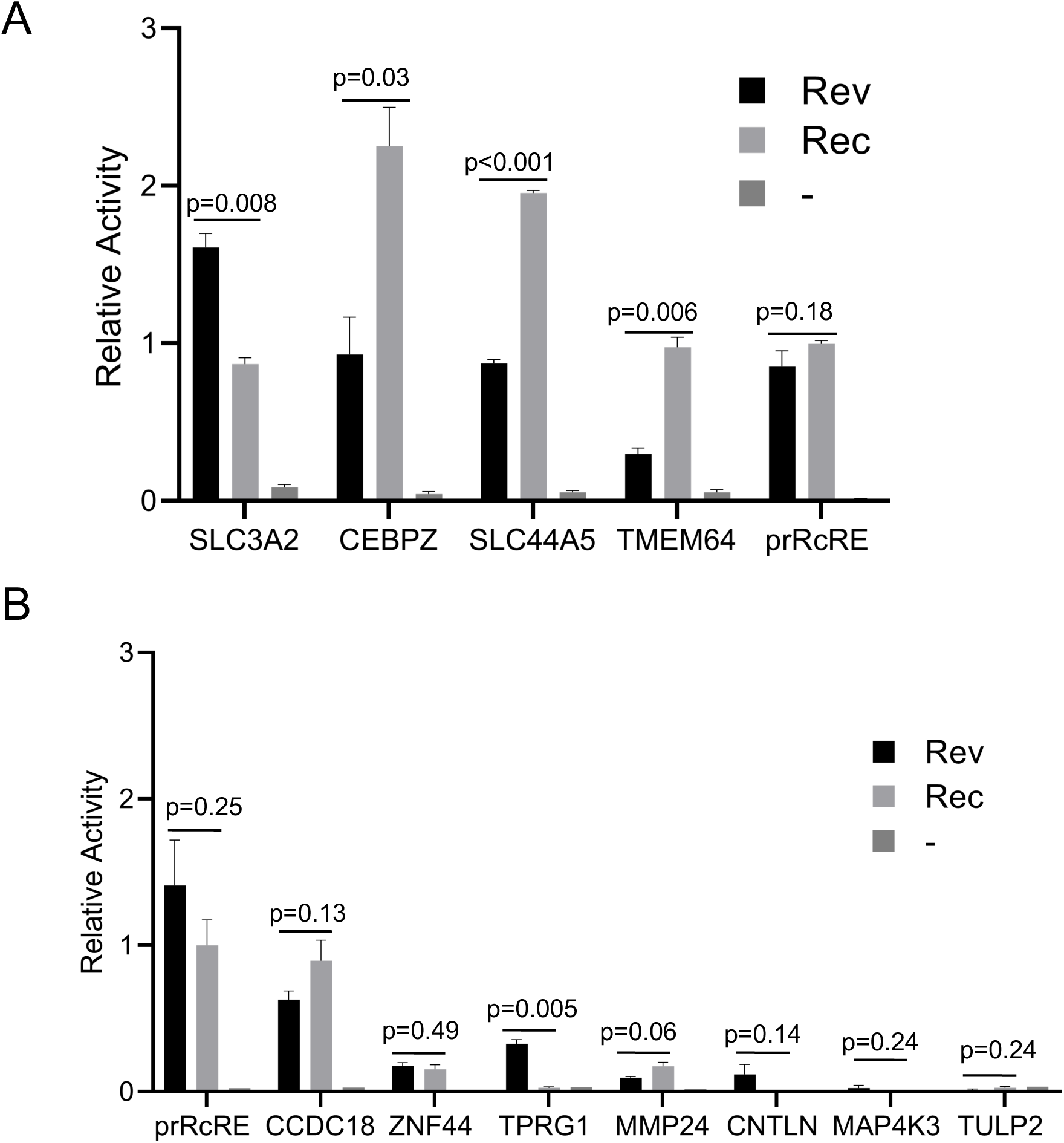
Functional activity of selected RNA elements in a transient transfection assay. The functional activities of eleven of the RcREs identified with the targeted bioinformatic approach were measured by transfecting the reporter constructs containing each RcRE into 293T/17 cells together with either 50ng of prRev or prRec or no additional vector. As controls, reporter constructs containing the prRcRE and HIV-1 RRE were also tested. In addition, a plasmid expressing Secreted Alkaline Phosphatase (SEAP) was transfected with each sample as a normalization control for transfection efficiency. After 72 hours, cell supernatants were harvested and supernatant p24 and SEAP were measured. The samples were then normalized for differences in transfection efficiency using the SEAP values. For comparison between experiments the SEAP normalized p24 value for the Rec/prRcRE pair was set to 1 and the activity of each Rev/RcRE or Rec/RcRE combination relative to this normalized value was plotted. The error bars show the variation between two independent transfections performed at the same time. To determine p values, between Rev and Rec, a two-tailed student t-test was performed using GraphPad Prism 8. The results shown in A and B were from transfections performed on different days. In panel A, the raw prRec/prRcRE p24 value before normalization was 6.4 ng/ml and in panel B it was 4.8 ng/ml.

We next tested seven of the ten additional potential RcRE elements (CCDC18, MMP24, TPRG1, CNTLN, ZNF44, TULP2 and MAP4K3) that were identified in the overlap of our RcRE database and the dataset of genes that were differently regulated after HIV infection. Two of the elements, MAP4K3 and TULP2, did not produce detectable levels of p24 above background with either Rev or Rec and MMP24 responded very weakly (Figure 4B). Four of the seven elements (CCDC18, TPRG1, ZNF44 and CNTLN), showed increased activity with Rev (Figure 4B), but only CCDC18 and ZNF44 functioned with Rec. Taken together, these results demonstrate that multiple RcREs within human genes can function with both Rev and Rec and that small sequence differences in the RcREs can result in significant functional differences and different activity with Rev and Rec. The fact that many functional RcREs were found in genes that were differentially expressed in HIV infected cells and that some of these functioned better with Rev than with Rec suggests that Rev could alter expression of the genes containing these RcREs after HIV infection.

### Functional testing of RcRE sequences with different HIV-1 Rev proteins

Rev sequences derived from primary isolates of subtypes A, G and the recombinant virus CRF02 AG were previously shown to have differential activity in conjunction with the pNL4-3 RRE. To test how some of the selected RcRE sequences would function with these different Rev proteins, eight HIV-1 Rev sequences were tested with reporter vectors containing the CEBPZ, SLC44A5, SLC3A2 and TMEM64 RcREs (39). All of the Rev proteins were expressed from pCMV vectors that were identical except for the Rev sequences. We also included the prRev and the prototype Rec protein in these experiments and a plasmid expressing SEAP was also added to each transfection as a normalization control.

As shown in Figure 5A all of the Rev proteins functioned on the prRcRE, however their activity varied greatly, and they functioned in a different rank order compared to their function on the NL4-3 RRE, as shown in our previously published report (39). They also demonstrated varying levels of activity on all of the RcREs (Figure 5B). In order to more easily compare how a particular Rev protein behaved with each of the RcREs, we normalized the activity of each Rev/prRcRE to 1 and plotted the activity of each Rev/RcRE combination relative to the normalized value (Figure 5C). In this way, it can be easily seen that some Rev proteins function more efficiently with certain RcREs. For example, 6AG Rev functioned better with the CEBPZ RcRE and less well with the TMEM64 RcRE compared to its activity in conjunction with the prRcRE, whereas 1A Rev functioned about the same on the CEBPZ RcRE and less well with the TMEM64 RcRE compared to its activity with the prRcRE. On the other hand, 8G Rev functioned less well with the CEBPZ RcRE, and hardly at all with the TMEM64 RcRE compared to its activity with the prRcRE. We then tested seven additional RcRE sequences (CCDC18, MMP24, TPRG1, CNTLN, ZNF44, TULP2 and MAP4K3) with three of the HIV-1 Revs (1A, 6AG and 8G) that showed high, medium and low p24 expression on the HERV-K prRcRE. Again, the different RcREs functioned with differential activity with the different Revs and Rec. However, in these cases, the rank order of activity of all of the RcREs tested remained nearly the same with all of the Rev proteins, with minor variations compared to Rec (Figure S1A and S1B). Taken together, these results indicate that different RcRE sequences respond differentially to different Rev proteins, suggesting that Rev proteins from different HIV primary isolates will likely influence host cell gene expression in different ways.

**Figure 5.**
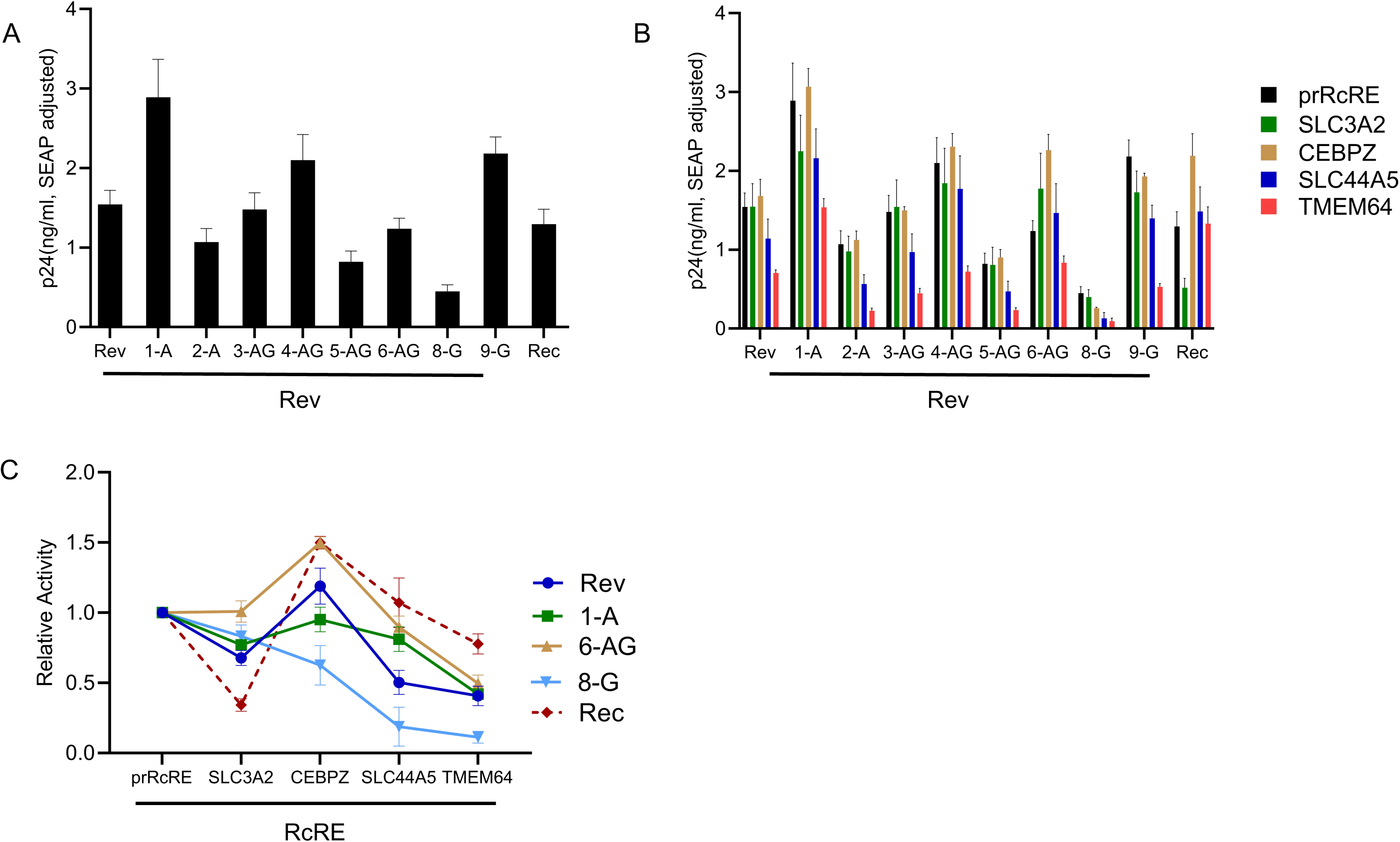
Functional activity of HERV-K RcRE sequences with HIV-1 Rev subtypes. The functional activity of four RcREs found within CEBPZ, SLC3A2, SLC44A5 and TMEM64 gene regions was measured by transfecting the reporter constructs containing each identified RcRE as well as the control prRcRE into 293T/17 cells together with either 50ng of prRev, prRec or one of eight different HIV-1 Revs (1A, 2A, 3AG, 4AG, 5AG, 6AG, 8G and 9G) In addition, a plasmid expressing Secreted Alkaline Phosphatase (SEAP) was transfected with each sample as a normalization control for transfection efficiency. After 72 hours, cell supernatants were harvested and supernatant p24 and SEAP were measured. The samples were then normalized for differences in transfection efficiency using the SEAP values. (A) Vectors expressing Rev from various HIV subtypes, prRev and prRec were transfected together with a reporter plasmid that contained the prRcRE. (B) Vectors expressing Rev from various HIV subtypes, prRev or prRec were transfected together with reporter plasmids that contained the prRcRE or the RcREs found within the CEBPZ, SLC3A2, SLC44A5 and TMEM64 gene regions. (C) For each of the Rev (or Rec) sequences, the SEAP adjusted p24 value of the Rev (or Rec)/ prRcRE pair from part B was set to 1, and the activity of that specific Rev (or Rec) with each of the other RcREs was plotted relative to this value. This allows a direct comparison of the activity of each Rev (or Rec) protein with each of the different RcREs relative to its activity on the prRcRE.

### Identification of HERV-K transcripts present in total and cytoplasmic RNA in cells transduced with retroviral vectors expressing HIV Rev and/or Tat or HERV-K Rec

To directly analyze if expression of regulatory HIV proteins (Tat and Rev) and the HERV-K Rec protein could change HERV-K mRNA expression in T-lymphoid cell lines, we transduced SupT1 cells with MSCV-derived retroviral vectors expressing Rev, Tat, Rec or both Rev and Tat. As a control, we also transduced cells with a retroviral vector lacking an insert, empty vector (EV). Each retroviral vector also expressed a fluorescent protein from an Internal Ribosome Entry Site (IRES) (see Methods) to allow identification and sorting of the cells that were transduced. Forty-eight hours post transfection, cells were sorted by flow cytometry to isolate the transduced populations. Both total and cytoplasmic polyA+ RNA was isolated and stranded cDNA libraries were prepared for RNA sequencing (RNAseq). Sequencing was performed on the Illumina HiSeq platform (see Methods). Trimmed reads were aligned to the hg19 version of the human genome using HISAT2. The BAM files were then filtered to obtain only the reads that mapped uniquely to one proviral location. To be able to count reads emanating from HERV-K proviruses, we created a Gene Transfer Format (GTF) file integrating all known HERV-K proviral genomic coordinates into the Ensembl GTF, hg19 build (44). This enabled us to identify the HERV-K loci that were transcriptionally active (14, 17).

When total RNA was analyzed, expression from 9 HERV-K loci (1q21.3, 3q12.3, 4p16.1b, 6p21.1, 6q25.1, 11q12.3, 19q13.12b, 20q11.22 and 22q11.23) was identified in all samples with only very slight differences between the samples transduced with the different vectors. Some reads also mapped to HERV-K locus 11q22.1, but only in the sample that received the Rec vector (Figure 6A). These reads appeared to originate from the exogenously expressed Rec, since the sequence of the reads were a perfect match to the sequences of Rec that are in the vector. When cytoplasmic RNA was analyzed a very different picture emerged. The samples that were expressing Rev or both Tat and Rev contained an increased number of reads for HERV-K proviruses 3q12.3, 4p16.1b and 22q11.23, with the greatest differences being seen for the 3q12.3 virus (Figure 6B).

**Figure 6.**
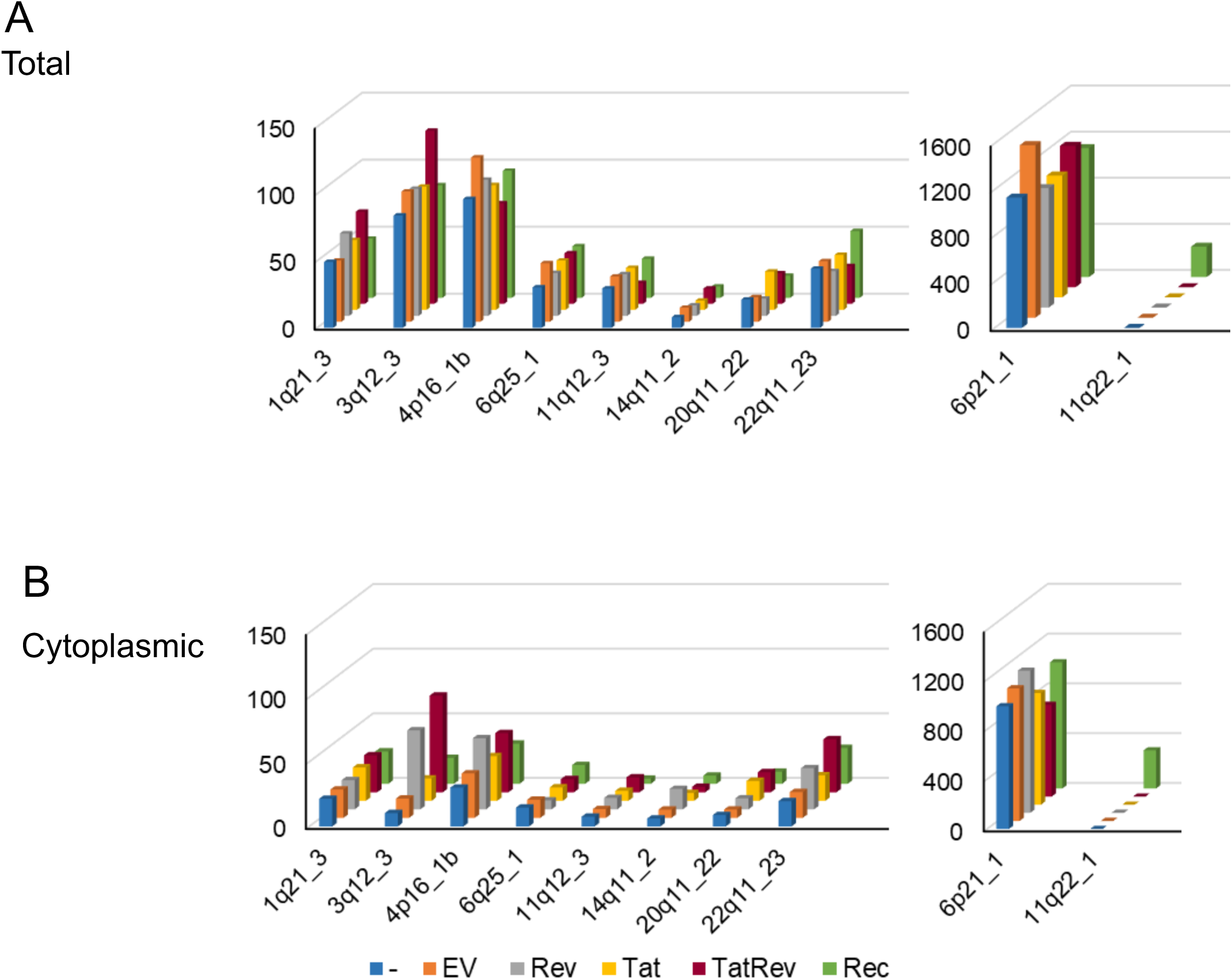
Identification of HERV-K proviral transcripts from total and cytoplasmic RNAseq data. SupT1 cells were transduced with retroviral vectors expressing a fluorescent protein marker and Rev, Tat, Rec, a combination of Rev and Tat, or only the fluorescent protein (EV). The fluorescent transduced cells were isolated by flow cytometry. SupT1 cells that were mock transduced and subjected to collection by flow cytometry were also included as a control (-). At 48hrs post transduction, both total and cytoplasmic RNA was prepared and stranded RNAseq libraries were constructed. Illumina sequencing was performed yielding 2x 150bp paired-end reads. Reads were then mapped to the human genome (hg19) using HISAT2 and HERV-K loci were quantified with DESeq2 using a custom GTF file. Normalized read counts that mapped uniquely to each HERV-K locus are shown for (A) total RNA or (B) cytoplasmic RNA. The reads mapping to 11q22.1 in the Rec sample are derived from the vector that expressed Rec and not the proviral locus. Quantitation of these reads and the reads mapping to 6p21_1 are presented in the right panel because they are expressed at a higher level than the others.

A quantitative analysis, using DESeq2, of the read counts mapping to 3q12.3, is presented in Figure 7. In this virus, the reads mapped across the entire proviral genome (Figure 7A). Cytoplasmic samples from cells transduced with a Rev-expressing construct, had 4.2 fold (*p* value = 1.13E-08) more reads uniquely mapping to this locus compared to the cells transduced with the empty vector and the cytoplasmic samples from cells transduced with both Tat and Rev had 5.4 fold (*p* value = 4.10E-10) fold more reads (Figure 7C). The fact that the increase in read counts maps across the entire proviral locus strongly suggests that Rev has facilitated the nucleocytoplasmic export of the full length unspliced HERV K genomic RNA in the transduced cells. Interestingly, cells transduced with HERV-K Rec, showed only a small increase in reads (1.4 fold). The differences in the number of reads mapping between total and cytoplasmic RNA samples in the other 2 proviruses were also quantified but were much less dramatic (1-2 fold). (Figure S2)

**Figure 7.**
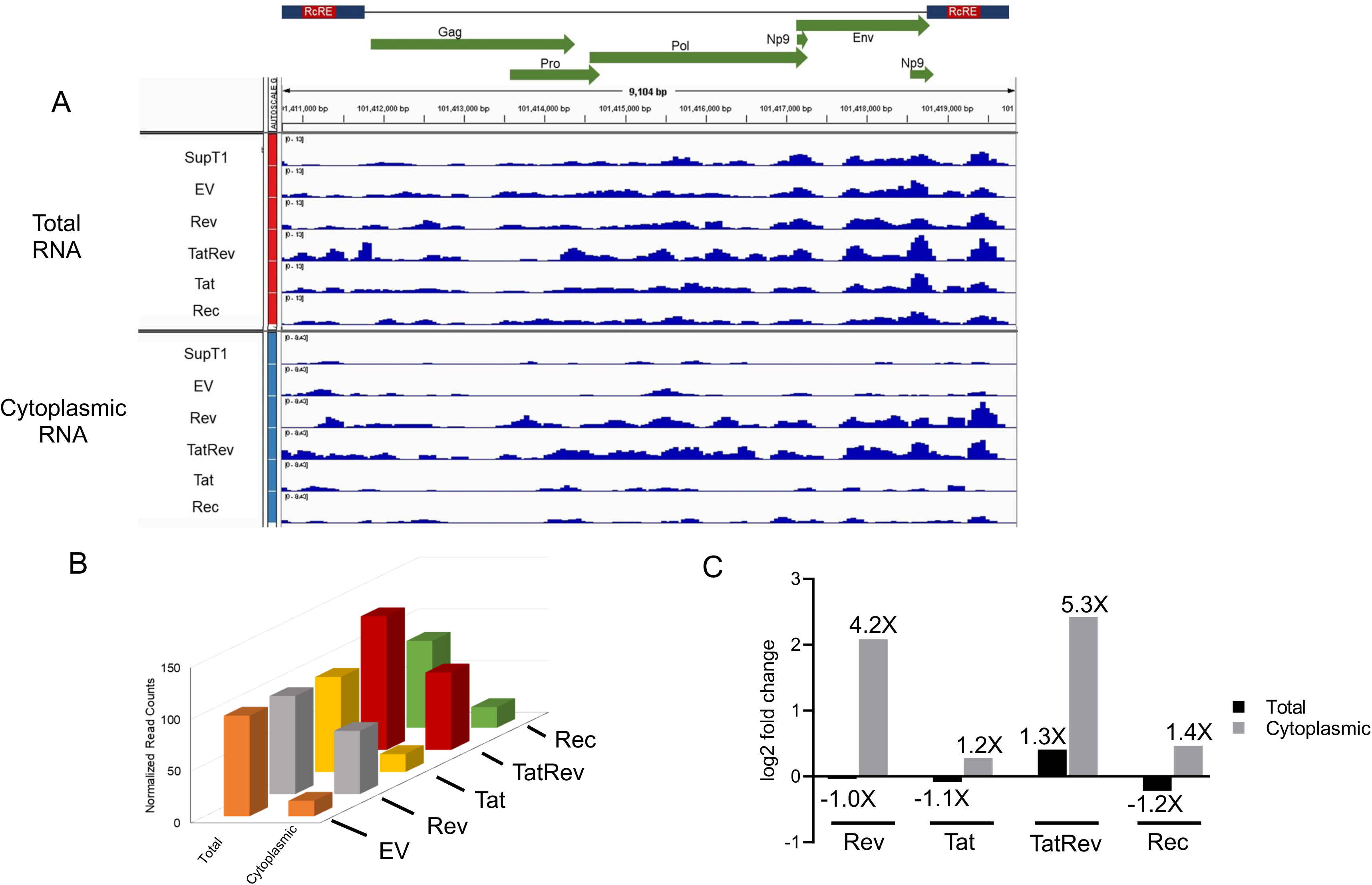
Quantitative analysis of HERV-K 3q12.3 proviral transcripts from total and cytoplasmic RNAseq data. (A) Total and cytoplasmic DESeq2 normalized read counts for HERV-K 3q12.3 provirus, were visualized with IGV. They are shown below a diagram of the genome of the provirus showing the open reading frames and LTRs which contain the RcREs. (B) The number of normalized reads mapping uniquely to the HERV-K 3q12.3 locus for total and cytoplasmic RNA are plotted for each of the samples transduced with either empty vector (EV), Rev, Tat, both Tat and Rev or Rec. (C) Quantitation of the data in (B) showing the fold change in read count relative to the sample transduced with the empty vector (EV).

As a control, we also examined the reads mapping to the gene encoding NEAT1 lncRNA. This gene is known to encode a full length unspliced RNA (NEAT1_2) that remains nuclear, localizing to paraspeckles, as well as a shorter form whose localization is less clear (NEAT1_1) (46). Figure S3 shows that reads derived from total RNA mapped across the whole NEAT1 gene region. However, in cytoplasmic RNA, only reads mapping to the region corresponding NEAT 1_1 were present. This serves as a control to confirm that our cytoplasmic RNA prep was not contaminated with RNA that should remain nuclear.

### The 3’ LTR of the HERV-K 3q12.3 provirus contains a RcRE that functions better with Rev than with Rec

Since transcribed HERV-K 3q12.3 mRNA was efficiently exported to the cytoplasm in the presence of Rev, it was of interest to analyze whether the 3q12.3 provirus contains RcRE sequences that function in conjunction with Rev. Since the RcRE is present in the U3 region of the HERV-K LTR, only the 3’ RcRE would appear in RNA transcribed from the provirus. Nevertheless, we tested both the 5’ and 3’ RcRE sequences in this provirus by cloning them into the Gag-Pol reporter construct and testing them with the different Rev proteins that we used in the experiments described above.

As shown in Figure 8A, all of the HIV-1 Rev proteins functioned well on the 3’ RcRE from this provirus. In contrast, very little function was seen in conjunction with HERV-K Rec. This correlated well with the RNAseq data that showed that only Rev significantly promoted the export of the HERV-K proviral RNA containing this RcRE. The different Rev proteins functioned in a somewhat different rank order than they did on the prRcRE, highlighting the fact that different Rev proteins show different activity on the RcREs due to unknown factors that affect their interactions. As in shown in Figure 8B, the 5’ RcRE from 3q12.3 was completely unresponsive to any of the proteins tested in spite of the fact the two RcREs are highly similar, showing 96% sequence identity to each other and 96.7% (3’) and 95.6% (5’) sequence identity to the prRcRE (Figure 8C).

**Figure 8.**
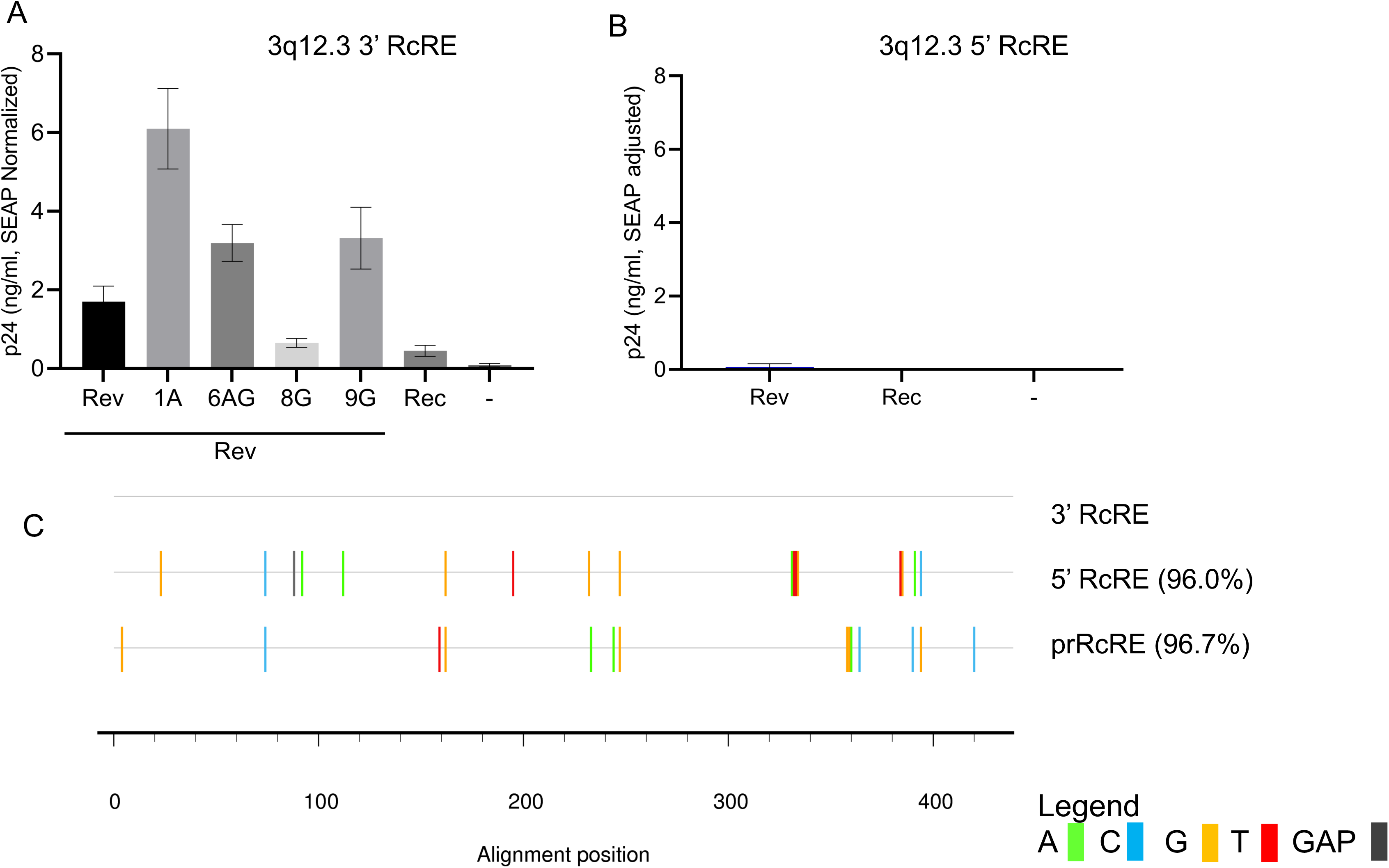
Functional activity of HERV-K 3q12.3 proviral RcREs in a transient transfection assay. Both the 5’ (A) and 3’ (B) RcRE sequences from HERV-K 3q12.3 were cloned into the Gag-Pol reporter construct which was then transfected into 293T/17 cells together with either 50ng of prRev, prRec or Rev from HIV-1 subtypes 1A, 6AG, 8G or 9G. In addition, a plasmid expressing Secreted Alkaline Phosphatase (SEAP) was transfected with each sample as a normalization control for transfection efficiency. After 72 hours, cell supernatants were harvested and supernatant p24 and SEAP were measured. The samples were then normalized for differences in transfection efficiency using the SEAP values. (C) A sequence alignment comparing the prRcRE and the 5’ RcRE of HERV-K 3q12.3 to the 3’RcRE of HERV-K 3q12.3. Nucleotide mismatches are shown by the colored lines. The 5’ RcRE and prRcRE were 96.0% and 96.7% identical to the 3’ RcRE.

## Discussion

Many studies have described increased HERV-K expression in HIV infection and HIV infected cells (33–36, 47). There have also been reports of immune responses to HERV-K in HIV infected individuals, and, in a recent report, it was shown that antibodies directed against HERV-K Gag were present in elite controllers (48). It has also been shown that HIV Tat can enhance HERV-K transcription from several proviral loci (37, 38). However, there have been no previous studies that have analyzed the effects of Rev expression on HERV-K expression, although studies conducted many years ago reported that Rev can enhance expression of unspliced RNA from reporter vectors through direct interactions with prototype RcREs cloned from unknown locations in the human genome (23, 31). Based on the advances in sequencing that led to the publishing of the human genome, we now know that there are several HERV-K proviruses that contain potential functional RcREs (14, 17). In addition to these, there are many solo HERV-K LTRs that are likely to contain functional RcREs (14).

Curiously, the previously studied functional RcREs do not show an exact sequence match to any RcREs identified in the current versions of the human genome. However, here we show that Rev can indeed function in conjunction with many of the HERV-K RcREs that are present in the human genome and that many of them are in protein-coding gene regions. Some of the RcREs that we identified and tested have the same sense as the coding region of the gene they reside in. However, several of the RcREs are in the anti-sense orientation relative to overlapping protein coding genes. In some cases, a non-coding potential antisense RNA has been identified with the same sense as the RcRE, in other cases nothing is known about transcription on the “anti-sense strand. Recently it has been shown that many genes encode overlapping anti-sense RNA (49, 50), which can sometimes change regulation of a protein coding gene on the opposite strand. Thus, it is possible that Rev interactions with RcREs in anti-sense RNAs could modulate gene expression (19). Whether this occurs in the case any of the genes where we have identified “anti-sense” RcREs (see Table 1) requires further study.

Most interestinglty, our studies clearly show that Rev can act to promote export of endogenously expressed HERV-K RNAs. The provirus that showed the most significant response to Rev was the provirus located at 3q12.3. This provirus has previously been shown to be transcriptionally active in many cells, including HIV infected cells (14, 51, 52). Our results showed that expression of Rev led to an increase of the full-length mRNA in cytoplasmic RNA and also that expression of Tat and Rev together led to an increase of this RNA in total RNA. This suggests that Tat increased transcription, generating mRNA that could be exported by Rev.

Our data also clearly show that different Rev proteins function with different efficiencies in conjunction with different RcREs and that some RcREs function better with some Rev proteins than with Rec. This result dovetails nicely with our previous finding that Rev proteins and RREs from different primary isolates can have significantly different levels of activity.

The fact that HIV-1 Rev promotes the nucleo-cytoplasmic export of unspliced mRNAs expressed from the Type I 3q12.3 HERV-K provirus, whereas Rec has little effect was an especially surprising result. Type 1 viruses cannot express Rec because of the deletion of essential parts of the Rec ORF. However, since Rec can act in *trans*, endogenous Rec expression could lead to export of unspliced mRNA from Type I proviruses, if a functioning RcRE is present. However, in the case of 3q12.3, our results demonstrated that both the 5’ and 3’ RcREs present in the LTRs of this provirus function very poorly with the prototypical Rec that was tested. Although it remains possible that different endogenous Rec proteins expressed in other cells may work better in conjunction with the 3q12.3 RcRE, it is clear that in SupT1 cells the unspliced mRNA expressed from this provirus is only exported to the cytoplasm in response to Rev expression. We do not know if there are other HERV-K proviruses that will respond to Rev and Rec in a similar fashion, since SupT1 cells only show RNA expression from a few. Since HERV-K proviruses and their expression are polymorphic in the human population and there are many different HIV subtypes, there is a clear potential for differential effects of HERV-K/HIV interactions in different infected individuals. This should be taken into account in studies of the potential role of HERV-K in cancer and other diseases in HIV patients, since the effects on HERV-K expression could be very different depending on both the endogenous HERV-K expression profile and the HIV sequence.

Previously published reports have examined HERV-K(HML-2) proviral expression in uninfected and HIV-1 infected primary CD4+ cells (51, 52). In these studies, it was found that two HERV-K loci, 1q22 and 3q12.3, produced the majority of the proviral HERV-K RNAs. Five additional loci, 1q23.3, 6q25.1, 8q24.3, 19q13.42 and 12q24.33, were also transcriptionally active, albeit at lower levels than 1q22 and 3q12.3. Another report studied HIV-1 infection in primary CD4+ T-cells and noted a small increase in overall HERV-K expression compared to uninfected cells (32). However, our further analysis of this RNAseq dataset (data not shown), reveal eight HERV-K loci, 1q21.3, 1q22, 1q23.3, 3q12.3, 4p16.1a, 11q12.3, 14q11.2 and 19q13.12b, as being transcriptionally active. Importantly, the RNA from the provirus at 3q12.3 that contains the RcRE that responds to Rev, was identified as active in all three previous studies using primary cells. Furthermore, in the HIV-1 infected CD4 T-cells, the RNA from 3q12.3, increased by 1.44 fold (pvalue:0.01) compared to uninfected CD4+ T-cells (32). RNA expression from two HERV-K loci, 1q22 and 1q23.3, were in common in the three published reports examining primary CD4+ T-cells, but were not seen in our analysis of RNA in SupT1 cells.

Whereas the effects of Rev on proviral HERV-K expression may be the most important finding in our study, we started by identifying potential RcREs in HERV-K solo LTRs present in non-viral protein coding and non-coding genes. We originally based our selection of sequences to test based simply on how similar they were in sequence to the prototype RcRE that was analyzed in previous studies. While this did indeed lead to the identification of RcREs that functioned with varying efficiencies with both Rev and Rec, it became clear that primary sequence is not a good indicator of RcRE function. This may be best illustrated by fact that the RcRE located within the 3’ LTR from the 3q12.3 HERV-K provirus was shown to function extremely well with some HIV Revs, but not with others and also rather poorly with Rec (see Figure 8). In contrast, the 5’ RcRE from this provirus did not function with either Rev or Rec, in spite of the fact that there were only a few primary sequence differences between the 5’ and 3’ RcRE and the prototype RcRE. Additionally, the RcRE which mapped to the SLC3A2 region clearly functioned better with the prototype Rev compared to its activity with the prototype Rec. These results highlight that functionality cannot currently be predicted simply by an RcRE sequence analysis. Thus, functional activity experiments using reporter assays like the one we describe here are essential to evaluate the extent to which a specific RcRE will function in conjunction with either a specific Rev or Rec sequence.

Our RNAseq analysis also identified two more HERV-K proviruses (4p16.1b and 22q11.23) where there was an increase in the amount of unspliced mRNA in the cytoplasm, when HIV-1 Rev or Tat and Rev were present, although to a much lesser extent than seen for 3q12.3. While 3q12.3 is a full-length intact provirus with a functional 3’ RcRE, the 4p16.1b and 22q11.23 proviruses have truncated 3’ LTRs with only partially intact RcREs. It remains to be determined how Rev may act to promote nucleo-cytoplasmic export of the mRNA expressed from these two proviruses.

Previously published reports demonstrate that transcription for both 3q12.3 and 4p16.1b is initiated in their respective 5’ LTRs, while the transcription for 22q11.23 is through a separate LTR, located a little over 500bp upstream of the provirus (53). Notably, it has been reported that the 5’ LTR from the provirus 3q12.3 has evolved to include a duplication of the HOX-PBX transcription binding site as well as a SNP in the RFX3 transcription factor binding site (54). This may explain why this provirus is highly transcribed in many different cell types and why it no longer is able to function as a RcRE.

Most reading frames in HERV-K 22q11.23 and 4p16.1b have premature stop codons, but although the *rec* gene of 4p16.1b is heavily mutated relative to the prototype HERV-K its reading frame remains open. The 3q12.3 provirus, on the other hand, has maintained open reading frames for both *gag* and *np9*. Thus, Rev mediated export of mRNA expressed from the 3q12.3 provirus could lead to the expression of HERV-K Gag and the formation of immature HERV-K virus particles. Whether this could occur with this or other HERV-K proviruses in an infected individual would depend on the combination of the RcRE that is present in a transcribed provirus and the particular Rev protein that was expressed. Consistent with this notion, an increase in HERV-K mRNAs and proteins, as well as antibody responses to HERV-K during HIV-1 infection, have previously been reported (36, 48, 55, 56). A recent article reporting on increased HERV-K capsid antibody levels in elite controllers suggest that immune responses to HERV-K structural proteins could contribute to HIV immune control (48). Thus it will be of clear interest to further define the RcREs present in HERV-K mRNAs that are expressed in HIV infected individuals, especially those with strong anti-HERV-K immune responses, and to determine whether the Rev proteins expressed in these individuals can function in conjunction with these RcREs. It has also been reported that the 3q12.3 provirus is actively transcribed in patients with Amyotropic Lateral Sclerosis (ALS) (57) and although this remains controversial, it has been proposed that HERVs plays a pathophysiological role in this lethal disease (for a review see (58). Some HIV-infected patients develop neurological manifestations that resemble classical ALS, although it occurs at a younger age and there has been reports of improvement following the initiation of antiretroviral therapy (59). In view of the results presented here, it may be of interest to address whether HIV Rev expression leads to increased expression of HERV-K Gag proteins in the brain and immune responses to HERV-K in these patients.

The mechanism(s) responsible for the difference in relative HIV-1 Rev or HERV-K Rec activity when paired with different HERV-K RcREs was not examined in this study. It has been previously proposed that different stem-loops in the prototypical HERV-K RcRE sequence may be involved in binding to Rev and Rec (31, 60, 61). For HIV-1 Rev, it has been suggested that there may be two discrete binding sites that lie within stem-loop 2 and 6, while for HERV-K Rec, binding between the protein and RNA may be much more complex, involving folding and three-dimensional structure rather than discrete binding sites within the HERV-K RcRE. Our results in this study demonstrate that the RcREs that are found within SLC3A2, TPRG1 and CNTLN displayed higher functional activity levels with HIV-1 Rev than with HERV-K Rec. Studying additional HERV-K RcREs in other areas of the genome and their functional activity as well as performing SHAPE analysis may be helpful in further pinpointing areas of the RcREs that are important for HIV-1 Rev binding.

## Conclusions

During this study we identified HERV-K RcRE sequences that functioned with varying efficiencies in response to different HIV-1 Rev proteins. Some of the HIV-1 Rev proteins functioned better with several of the selected HERV-K RcREs than with HERV-K Rec. Additionally, we showed that the RcRE found within the 3’ LTR of HERV-K 3q12.3 functioned well with many different Rev proteins, but hardly at all with the prototypical Rec protein. Consistent with this, we showed that expression of Rev in SupT1 cells resulted in a significant increase in cytoplasmic, unspliced viral mRNA from the HERV-K 3q12.3 provirus which has a complete open *gag* reading frame. HIV Rev could thus change HERV-K protein expression, resulting in novel immune responses to these viruses. Finally, our results indicate that effects of HIV Rev on HERV-K expression in infected individuals will differ depending on variation in HERV-K expression and the HIV subtype.

## Methods

### Identification of genomic HERV-K RcRE sequences

A list of sequences highly similar to the HERV-K RcRE was generated through a BLAST search using the prototypical HERV-K RcRE sequence (23) to search the Ensembl DNA, non-coding RNA, and cDNA human genomic databases (GRCh38.87) (44). Sequences with greater than 94% sequence identity were identified and listed.

Two other gene lists were retrieved from previously published studies identifying differential gene expression during HIV-1 infection (32) and differential gene expression when HERV-K Rec was expressed in hECC (62). Genes were considered differentially expressed if the false discovery rate (FDR) adjusted p-value was p<0.01.

### Plasmids

All plasmids utilized were given the name pHRXXXX for identification and to facilitate requests. Synthetic double stranded DNA sequences (gBlocks™; Integrated DNA Technologies, Coralville, IA) corresponding to RcRE sequences from the gene regions identified in the gene list were synthesized and cloned into pHR2739, a pCMV-GagPol reporter plasmid lacking the RRE (63) such that the RcREs were positioned immediately downstream from the intron containing the GagPol coding region. To facilitate cloning, the gBlocks™ DNA fragments were designed with 25bp on either end overlapping the ends of the linearized pCMV-Gag-Pol plasmid. Fragments were cloned using the NEBuilder HiFi DNA assembly kit with a vector to insert ratio of 1:2, following the manufacturer’s directions. This created the plasmids pHR5342 (C11orf72), pHR5336 (cFLAR), pHR5341(U9linc), pHR5340 (RP4linc), pHR5534 (CEBPZ), pHR5540 (SLC3A2), pHR5538 (SLC44A5), pHR5536 (TMEM64), pHR5843 (CNTLN), pHR5908 (ZNF44), pHR5846 (CCDC18), pHR5684 (MAP4K3), pHR5835 (TPRG1), pHR5639 (TULP2), pHR5845 (MMP24), pHR5812 (3q12.3_3’_RcRE), and pHR5813 (3q12.3_5’_RcRE). pCMV-GagPol-RRE (pHR3442) has been used previously to measure Rev function (43). It was constructed in the same way as the pCMV-GagPol-RcRE plasmids described above, except that a DNA fragment containing the RRE, amplified by PCR from pNL4-3, was inserted into the backbone by traditional cloning methods rather than using the NEBuilder HiFi DNA assembly kit. The sequence of each inserted gBlock or PCR fragment in each plasmid was confirmed using Sanger DNA sequencing. Plasmids expressing Rev from several different HIV subtypes and the viruses from which they are derived have been previously described (40, 44). These included pHR5209 (subtype A-1), pHR5182(subtype A-2), pHR5183(subtype AG-3), pHR5185(subtype AG-4), pHR5187(subtype G-9), pHR5202(subtype AG-5), pHR5203(subtype AG-6), pHR5204(subtype G-8)). The plasmid expressing the prototypical Rev sequence (prRev), pHR30, which was derived from a clade B isolate, has been described previously (64).

### Cell Culture and Transient Transfections

Twenty-four hours prior to transfection, 1.7 x 10^5^ 293T cells were seeded into 12-well tissue culture plates in 1ml medium (DMEM, 10% Bovine Calf Serum (BCS), 50mg/ml gentamicin). To test the activity of the HIV-1 RRE and HERV-K RcREs, each well was transfected with 50ng of either pCMVRev (pHR30) or pCMVRec (pHR5313), 2µg of pCMV-Gag-Pol-RRE(pHR3442) or pCMV-Gag-Pol-RcRE(pHR5338) and 100ng of a plasmid expressing secreted placental alkaline phosphatase (SEAP) (pHR1831) using Turbofect™ transfection reagent (ThermoFisher). Cell supernatants were collected 72 hours post-transfection, and levels of p24 were determined by ELISA as previously described (65) The SEAP levels of the supernatants were determined by a chemiluminescent method (Roche) and were used to normalize the differences in transfection efficiencies.

### Retroviral vectors, production and transduction

The MSCV retroviral vector backbone was utilized to make vectors that express Rev, Rec or Tat as well as a fluorescent protein. Specifically, sequences encoding the respective proteins were cloned into an MSCV-derived construct, upstream of an IRES driving the expression of a fluorescent protein to create the constructs MSCV-Rev-IRES-eGFP (pHR5366), MSCV-Tat-IRES-mCherry (pHR5447) and MSCV-Rec-IRES-eGFP (pHR5450). MSCV-IRES-eGFP (pHR53650) was also utilized as an “empty” vector control (EV). The backbone pMSCV-IRES-fluorescent protein vectors were gifts from Dr. Dario Vignali (unpublished), obtained through Addgene.

For retroviral vector production, twenty-four hours prior to transfection, 8 x 10^6^ 293T cells were seeded into 15cm tissue culture plates in 20ml of growth medium (DMEM, 10% BCS, 50/ml gentamicin). Transient transfections were then performed using the calcium phosphate method (66) to deliver 40ug of each MSCV, 10ug of pMD2.G expressing VSV-G, a gift from Dr. Didier Trono (67) and 30ug of pHIT60 expressing MLV Gag-Pol, a gift from Dr. Alan Kingsman, Oxford University, Oxford, England (68) to each plate. After 5 hours, cell culture medium from the transient transfections was removed and replaced with 12ml of fresh medium. Viral vector stocks were harvested at 24, 48 and 72 hours then concentrated by ultracentrifugation (Beckman Coulter Optima XL-100) for 1 hour at 25K in an SW28 rotor and resuspended in PBS.

Vector titrations were performed by preparing six 2-fold dilutions of concentrated vector stocks (1:200, 400, 800, 1600, 3200, 6400, 12800) in PBS. One milliliter of each diluted stock was then added to 2.5 x 10^5^ SupT1 cells with 10ug of polybrene, incubated for 4 hours at 37°C in an incubator containing 5% CO_2_. The cells were then washed 2x with PBS and resuspended in 1ml of medium. After 48 hours, cells were harvested and washed 2X with PBS containing 10% BCS before they were analyzed for GFP or mCherry positive cells using the Attune NxT flow cytometer (Invitrogen). Virus titers were calculated when positive cells fell between 2-20% in the appropriate dilution using the following formula: (F x cells/volume) x dilution factor. Where F is the frequency of positive cells, cells is the total number of target cells infected, V is the volume of inoculum and the dilution factor is the virus dilution factor.

For cell transduction, SupT1 cells were split 24 hours prior to adding viral stocks to 1×10^6^ cells resuspended in 1ml of PBS at an MOI of 2 with 10ug/ml polybrene. Cells and virus were incubated for 4 hours at 37°C, 5% CO_2_, then washed twice and resuspended in 5ml of growth medium (RPMI, 10% FCS, gentamicin). After 48 hours, cells were sorted by flow cytometry for expression of GFP, mCherry, GFP and mCherry using the FACS Aria (BD Biosciences). Only cells expressing the appropriate fluorescent markers were chosen for RNA isolation.

### RNA isolation and sequencing

Total and cytoplasmic RNA from three biological replicates of SupT1 cells transduced with the empty MSCV vector (EV) or MSCV vectors expressing Rev, Tat, Tat and Rev and Rec together with the fluorescent proteins, were isolated from cells sorted by flow cytometry. The methods used for total and cytoplasmic RNA extraction were previously described (69), with the following modifications. For the total RNA extraction, the sorted SupT1 cells were pelleted, then washed twice with 5ml of cold phosphate-buffered saline (PBS). After the final wash, the cells were lysed in 500ul of lysis buffer containing 0.2 M Tris-HCL (pH 7.5), 0.2M NaCl, 1.5mM MgCl_2_, 2% sodium dodecyl sulfate (SDS) and 200µg of proteinase K per ml, and incubated for 1 to 2 hours at 45°C. For cytoplasmic RNA extraction, the sorted SupT1 cells were pelleted then washed twice with 5ml of ice-cold phosphate-buffered saline (PBS). After the final wash, the cells were pelleted and resuspended in reticulocyte standard buffer (RSB) (10mM Tris-HCL (pH 7.4), 10mM NaCl, 1.5mM MgCl_2_) followed by the addition of an equal volume of RSB with 0.1% IgePal™. Nuclei were cleared by two sequential centrifugations at 13,000 rpm in an Eppendorf centrifuge at +4C, prior to the addition of 2X PK buffer (200mM Tris-HCL (pH7.5), 25mM EDTA, 300mM NaCl, 2% SDS, 400µg of proteinase K per ml) to the cleared supernatant. The lysates were then incubated at 37°C for 30 minutes.

Total and cytoplasmic RNA lysates were then extracted once with an equal volume (1:1) of phenol and chloroform-isoamyl alcohol (24:1), followed by two extractions with only chloroform-isoamyl alcohol (24:1). Sodium acetate (3M, pH 5.5) was then added to the final extractions for a final concentration of 0.3M prior to precipitation with 2.5 volumes of 200 proof ethanol (−20°C). The RNA was pelleted for one hour at 4°C 3000 RPM (3014g) in a Baxter Cryofuge 6000 centrifuge, then washed twice with 70% ethanol and air dried prior to the addition of RNase/DNase free water. Total RNA was DNase digested (RQ1 RNase-free DNase, Promega) then extracted and precipitated as before. Final RNA concentrations were determined with a Qubit RNA HS assay kit (Qubit 3.0 fluorometer, ThermoFisher).

### NGS Sequencing

Stranded libraries were prepared by Novogene (Beijing, China) from total and cytoplasmic RNA using reagents from New England Biolabs (NEB). First, polyA+ RNA was isolated from total or cytoplasmic RNA using poly-T magnetic beads. The polyA containing RNA was then fragmented with fragmentation buffer (NEB)and first strand cDNA was synthesized using random hexamers and M-MuLV Reverse Transcriptase (RNase H-). Second strand cDNA synthesis was performed using DNA Polymerase I and RNase H while substituting dTTP with dUTP. The double stranded cDNA was purified using AMPure XP beads (Beckman Coulter, Beverly, USA). To ensure that the library was stranded, the second strand cDNA was digested by USER (Uracil-Specific Excision Reagent, NEB) prior to selecting for fragments approximately 150-200bp in length with AMPure XP beads. The final library was amplified by PCR, then re-purified with AMPure XP beads prior to sequencing on the Illumina platform at Novogene.

### Bioinformatics analysis

FASTQ files were trimmed with Trimmomatic-0.38 using the standard settings (ILLUMINACLIP:TruSeq3-PE.fa:2:30:10 LEADING:3 TRAILING:3 SLIDINGWINDOW:4:15 MINLEN:36), then assessed for quality using FASTQC software. Trimmed reads were aligned to hg19 (GRCh37) using HISAT2 (70) (hisat2 -p2 -q -x -1 $read1 -2 $read2) then passed through samtools view (-bS) to generate a .bam file for each alignment. The aligned reads were filtered to ensure that paired reads were uniquely aligned to the best sequence match one time using the following command: samtools view -h -f 0×2 file.bam | awk ‘substr($1, 0, 1)==”@” || $0 !∼ /ZS:/’ | samtools view -h -b > filtered_file.bam. A custom GTF file of the 94 HERV-K proviral chromosomal locations was created and incorporated into the GRCh37 GTF file to quantify HERV-K proviral aligned reads using Rsubread and FeatureCounts (71, 72). A new GTF file with the custom HERV-K GTF file incorporated into the GRCh37 GTF file was generated by merging the two files and then sorting on column 1 (chromosome number) followed by sorting on column 4 (chromosome start location). Differential expression analysis was performed using DESeq2 (73) on the reads quantified by featureCounts (71, 72) using the merged GTF file. Aligned reads for HERV-K 3q12.3 were visualized with IGV viewer after normalizing using DeepTools (74). BAM files for each condition were normalized using the bamCoverage argument (bamCoverage –bam yourfile.bam -o yourfile.norm.bw \ --normalizeUsing FPKM). The normalized bigwig files were merged with IGV tools to visualize the aligned reads with IGV for all replicates of each condition (74–76).

## List of Abbreviations

HERV: human endogenous retrovirus
HML-2: human MMTV-like
LTR: long terminal repeat
ORF: open reading frame
prRcRE: prototype RcRE
prRec: prototype Rec
EV: empty vector
IRES: internal ribosome entry site
SEAP: secreted placental alkaline phosphatase
GTF: gene transfer format

## Declarations

### Ethics approval and consent to participate

Not applicable

### Consent for publication

Not applicable

### Availability of data and materials

The datasets used and/or analyzed during the current study are available from the corresponding author on reasonable request.

### Author’s contributions

LRG wrote the first draft of the manuscript, prepared all figures and performed most of the experiments. REJ performed the functional experiments relating to HERV-K 3q12.3. PEHJ created vectors expressing the different HIV-1 Rev subtypes. SB provided bioinformatics expertise. MLH and DR conceived and contributed to the study design, manuscript revisions and project oversight. All authors reviewed and approved the final manuscript.

## Acknowledgements

We thank Jing Huang for expert technical expertise, David Grabski and Sarah French for helpful discussions and Dr. Dario Vignali for supplying the pMSCV-IRES-fluorescent protein vectors through Addgene. Technical assistance was provided by the University of Virginia Flow Cytometry Core Facility.

---

**Figure S1. Functional activity of HERV-K RcRE sequences with HIV-1 Revs.**

The functional activity of seven HERV-K RcREs was determined by transfecting the GagPol reporter constructs containing each identified RcREs as well as the control prRcRE into 293T/17 cells together with either 50ng of prRev, prRec or one of three different HIV-1 Revs (1A, 6AG and 8G). After 72 hours, cell supernatants were harvested and supernatant p24 was measured by ELISA. (A) Vectors expressing Rev from HIV 1A, 6AG and 8G, prRev or Rec were transfected together with reporter plasmids that contained the prRcRE or the RcREs from the seven indicated gene regions. Supernatant p24 was measured at 72 hours post transfection. (B) For each of the Rev (or Rec) sequences, the activity of the Rev (or Rec)/ prRcRE pair from part A was normalized to 1, and the activity of that specific Rev with each of the other RcREs was plotted relative to this value. This allows a direct comparison of the activity of each Rev (or Rec) protein with each of the different RcREs relative to its activity on the prRcRE.

**Figure S2. Quantitative analysis of HERV-K proviral transcripts from total and cytoplasmic RNAseq data.**

After normalization of the data, the fold difference in the number of unique reads mapping to the 22q11.23 (left panel) or 4p16.1b (right panel) loci were quantified using DESeq2 for total or cytoplasmic RNA from the Rev, Tat, Tat and Rev or Rec transduced samples, compared to the samples transduced with the empty vector.

**Figure S3**

**Visualization of NEAT1 reads from total and cytoplasmic RNAseq data.** Total and cytoplasmic DESeq2 normalized read counts for NEAT1 were visualized with IGV. Note that DNA reads from total RNA map across the entire NEAT1 gene region and include both the NEAT 1_1 and NEAT 1_2 RNA isoforms. In contrast, most of the reads from cytoplasmic RNA map only to the region corresponding to the NEAT1_1 RNA isoform.

