## Supplemental Figures for "HIV-1 Rev interacts with HERV-K RcREs present in the human genome and promotes export of unspliced HERV-K proviral RNA"

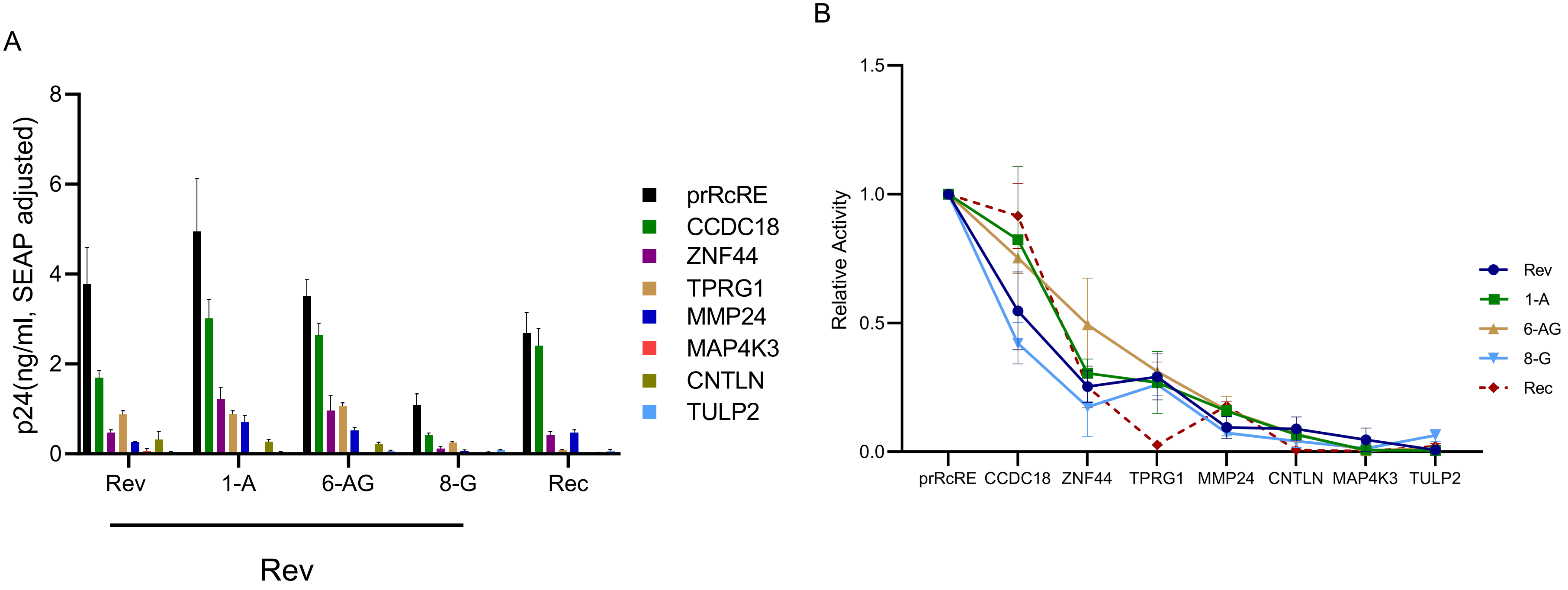


**Figure S1. Functional activity of HERV-K RcRE sequences with HIV-1 Rev subtypes.** The functional activity of seven HERV-K RcREs was determined by transfecting the GagPol reporter constructs containing each identified RcREs as well as the control prRcRE into 293T/17 cells together with either 50ng of prRev, prRec or one of three different HIV-1 Rev subtypes (1A, 6AG and 8G). After 72 hours, cell supernatants were harvested and supernatant p24 was measured by ELISA. (A) Vectors expressing Rev from HIV subtypes 1A, 6AG and 8G, prRev or Rec were transfected together with reporter plasmids that contained the prRcRE or the RcREs from the seven indicated gene regions. Supernatant p24 was measured at 72 hours post transfection. (B) For each of the Rev (or Rec) sequences, the activity of the Rev (or Rec)/ prRcRE pair from part A was normalized to 1, and the activity of that specific Rev with each of the other RcREs was plotted relative to this value. This allows a direct comparison of the activity of each Rev (or Rec) protein with each of the different RcREs relative to its activity on the prRcRE.

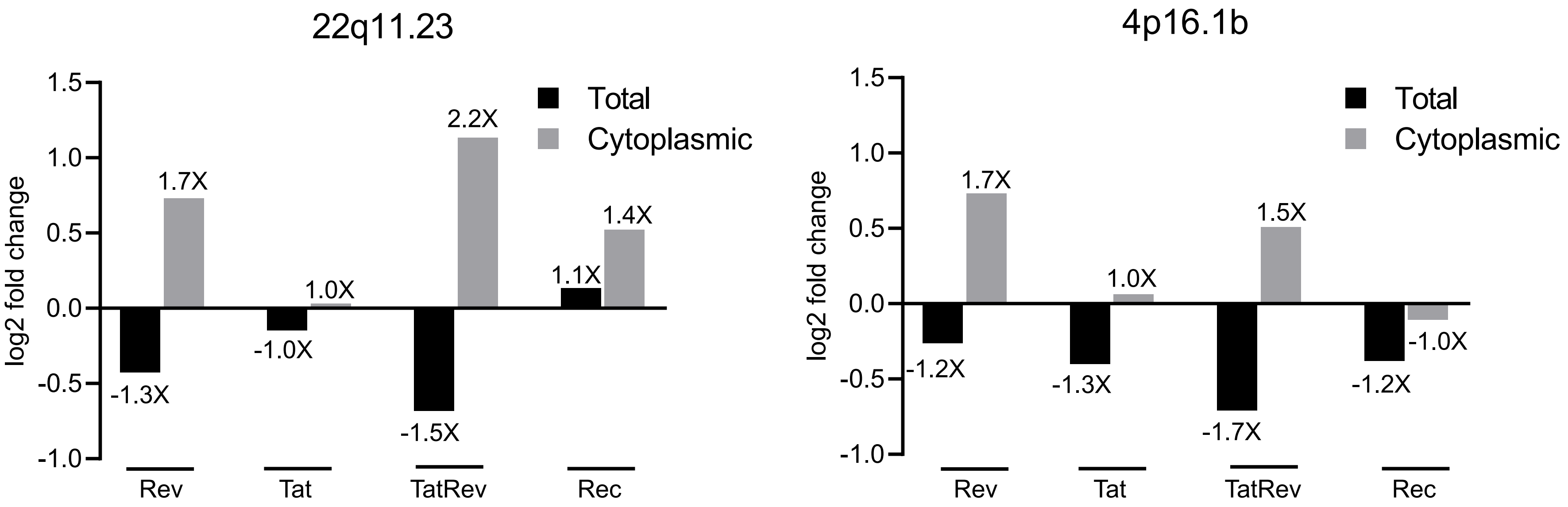


**Figure S2. Quantitative analysis of HERV-K proviral transcripts from total and cytoplasmic RNAseq data.** After normalization of the data, the fold difference in the number of unique reads mapping to the 22q11.23 (left panel) or 4p16.1b (right panel) loci were quantified using DESeq2 for total or cytoplasmic RNA from the Rev, Tat, Tat and Rev or Rec transduced samples, compared to the samples transduced with the empty vector.

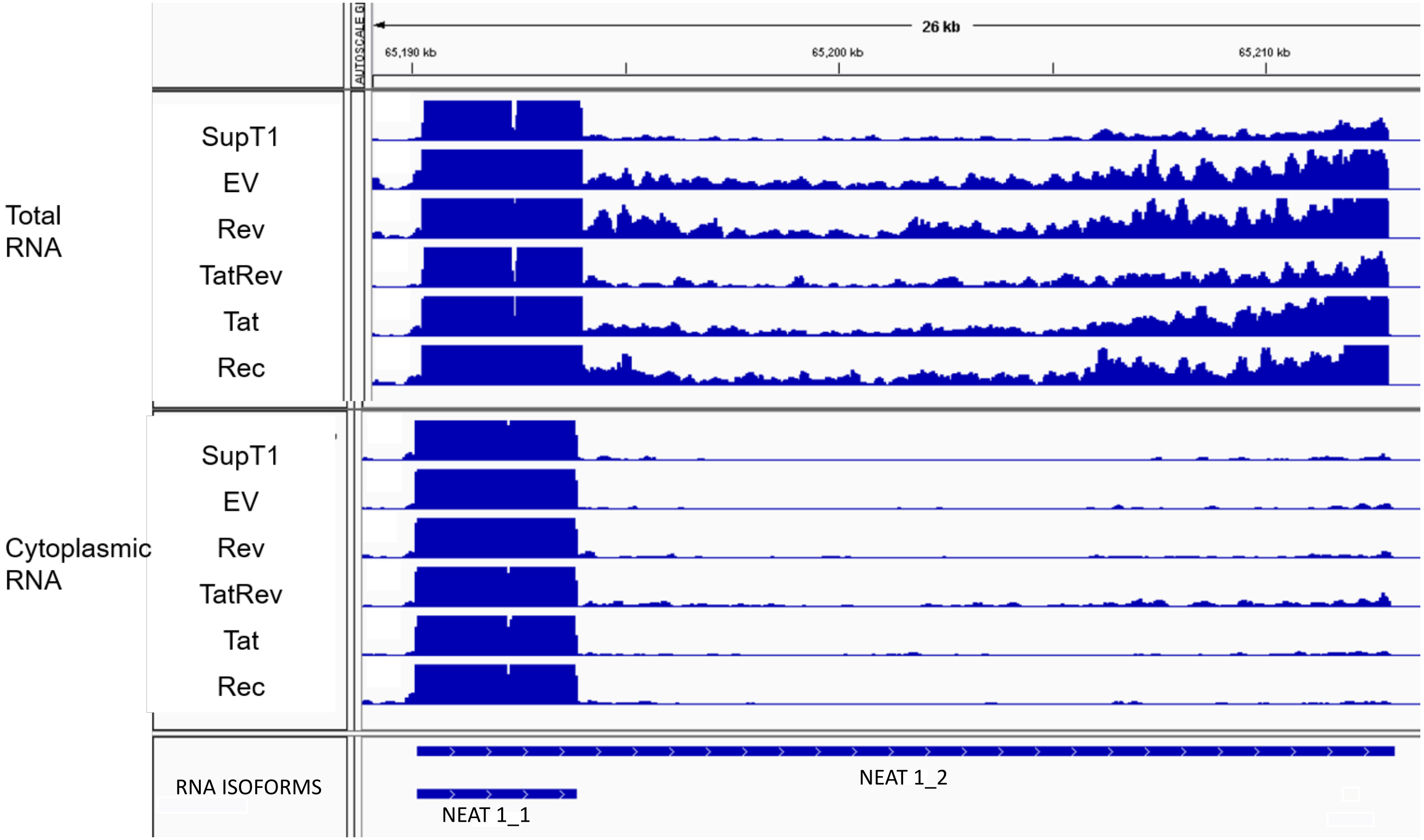


**Figure S3. Visualization of NEAT1 reads from total and cytoplasmic RNAseq data.** Total and cytoplasmic DESeq2 normalized read counts for NEAT1 were visualized with IGV. Note that DNA reads from total RNA map across the entire NEAT1 gene region and include both the NEAT 1_1 and NEAT 1_2 RNA isoforms. In contrast, most of the reads from cytoplasmic RNA map only to the region corresponding to the NEAT1_1 RNA isoform.
